# Raman Spectroscopic Study of the Influence of Oxytocin and *Uvariodendron anisatum Verdeck (Annonaceae)* Freeze-dried Extracts on Diet Induced Obesity in Sprague Dawley Rats

**DOI:** 10.1101/554956

**Authors:** Zephania Birech, Prabjot K. Sehmi, Peter W. Mwangi, Nelly M. Nyaga

## Abstract

Obesity is a condition affecting many people in the world. Obese people have increased risks of developing chronic metabolic diseases such as type II diabetes, hypertension, cancer among others. Early and rapid diagnosis of the condition together with effective treatment is therefore necessary. This work investigated, first, Raman spectroscopic similarities between oxytocin and a freeze-dried extract of a local herbal plant exhibiting oxytocin-like properties called *Uvariodendron anisatum Verdeck (Annonaceae)* (UAV). Secondly, whether Raman spectroscopy could be used for comparative studies of the influence of oxytocin and UAV on diet induced obesity in Sprague Dawley (SD) rat models. We also sought for obesity or metabolic syndrome biomarker Raman spectral bands. Both oxytocin and extract samples together with blood extracted from the rats were excited using a 785 nm laser with conductive silver paste smeared glass slides as Raman sample substrates. It was found that Raman spectral profiles of oxytocin solution and UAV freeze dried extract’s powder were identical with a cosine similarity value of 0.95 implying presence of similar Raman active molecules. The prominent peaks were those assigned to disulphide S-S stretching mode at 508 cm^−1^ and to tyrosine at 645 cm^−1^, 846 cm^−1^ and 1617 cm^−1^. Raman spectra of blood from oxytocin- and UAV-treated rats displayed similar profiles which were different from those of obese and non-obese (normal controls) animals. A prominent peak in spectra of treated rats centred at 401 cm^−1^ could be used as oxytocin biomarker band in blood. Comparison of average intensity trend of fructose bands at around 638 cm^−1^ and 812 cm^−1^ between prepared fructose solution and blood of treated rats, revealed elevated levels of fructose in blood of rats intraperitoneally injected oxytocin and UAV extracts. The result implied upregulation of fructose in oxytocin and UAV treated animals. Principal component analysis (PCA) showed that the Raman spectral profiles from blood of obese rats were different from those of non-obese rats. It also showed that spectra from oxytocin treated and UAV treated rat’s blood were similar indicating identical influence. The study shows the potential of Raman spectroscopy as tool for quick obesity (metabolic syndrome) screening with intensity of Raman bands associated with fructose acting as biomarkers. The same bands can also be used in comparative efficacy studies of anti-obesity drugs. Further studies are needed to validate these Raman spectroscopic results since, to the best of our knowledge, this was the first such investigation regarding comparison of UAV and conventional oxytocin together with their influence on obese SD rats. Weather human subjects exhibits similar results are also not known.

## 1. Introduction

Obesity, a metabolic condition characterized by abnormal increase in body weight and fat accumulation [1–3], is now a problem globally. According to World Health Organization (WHO), it was estimated that by year 2016 about 650 million people worldwide were obese [2]. The condition is caused by overconsumption of energy dense foods followed by less physical activity. There is a close relationship between being overweight and being obese. The two i.e. obesity and overweight are distinguished by a value known as body mass index (BMI) which basically is a ratio of weight (in kilograms) to the square of height (meters squared). An overweight and an obese human has a BMI value equal or greater than 25 and 30 respectively [2]. An obese person, therefore, is overweight. In rodent models, there is no universally agreed method of determining obese from non-obese rats but often those with fasting blood glucose (FBG) levels above 7 mmol/L [4] and those with increased volumes of subcutaneous and visceral adipose tissues [3] are regarded as obese. An obese individual has risks of developing chronic metabolic diseases such as type II diabetes, hypertension, coronary heart disease, cancer among others[1,3,5]. Management of this metabolic condition involves use of anti-obesity drugs, increase in physical exercise, reduction of high energy diet. These methods are un-popular due to side-effects and high failure rates. New interventions involving natural products with few side effects along with quick diagnostic techniques for monitoring their efficacy and at the same time detecting potential development of the condition are necessary.

One of the non-conventional potential alternative obesity treatments gaining a lot of attention lately involves use of oxytocin[6–8]. Oxytocin (OT) is a hormone associated with labor, lactation [9–11] and regulation of social behaviour in mammals [10,12]. It has chemical formula C_43_H_66_N_12_S_2_ and is locally produced in the brain and released to the circulatory system[12,13]. The compound consists of nine amino acids in the sequence: cysteine - tyrosine - isoleucine - glutamine - asparagine - cysteine - proline - leucine - glycine (CYIQNCPLG)[14]. The role of OT in weight reduction in obese rhesus monkeys[6] and in rats[7,15] has been reported. In mice[16] and male humans[16], OT caused a decrease in calorie intake. It has also been found that the hormone suppresses eating behaviour resulting in reduction of blood glucose levels, increase in insulin levels [10,17,18], reduction of glucose intolerance and insulin resistance[7] and a shift in diet preference from carbohydrates to fats[16]. The hormone is also reported to make cells resistant to diabetic conditions [13] and improve their insulin sensitivity [18]. In a study on African American males, the levels of oxytocin in blood of type II diabetic subjects were found to be lower than in healthy ones[19]. In the same study, subjects with higher levels of OT had lower body weights. Administration of OT is through intranasal [12,16], intraperitoneal [20] and subcutaneous[16] routes. Oral administration is rare due to its impaired and unpredictable absorption rates in the gastric system [21] though a review of it is being suggested elsewhere[22].

All these findings indicate a special role OT plays in the treatment and prevention of obesity and diabetes. Extended studies are, therefore, needed to investigate the influence of OT on metabolic diseases and other potential uses in non-conventional treatments.

In some parts of remote rural Kenya, pastoralists administer orally the herb *Uvariodendron anisatum Verdeck (Annonaceae)* (UAV) to cows that reject their calves soon after calving. The possible physiological explanation is that the herb stimulates the production of the bonding hormone oxytocin. This herb (i.e. UAV) is also used by traditional herbalists in parts where hospital facilities are distant to induce labor just as oxytocin does[23,24]. This work sought to investigate the similarities of UAV freeze dried extracts and oxytocin using Raman spectroscopy together with their influence on diet induced obesity in Sprague Dawley (SD) rats. This study, to the best of our knowledge, was the first of its kind. It was found that little Raman spectral differences exist between oxytocin and UAV extracts and no distinguishable differences were observed on their influence on obese SD rats. Administration of these two compounds (i.e. oxytocin and UAV extracts) via intraperitoneal injection to SD rats resulted in elevated levels of fructose in blood as revealed by intensity analysis of assigned Raman bands. Administration of these two compounds (i.e. oxytocin and UAV herbal extract) was done through intraperitoneal routes since this was the most viable option for the SD rats besides being the method with a higher clinical replicability.

## Materials and Methods

### Plant collection and Extract preparation

Fresh whole plants of *Uvariodendron anisatum Verdeck (Annonaceae)* collected from their natural habitat in Machakos County, Kenya. The identity of the plant samples confirmed at the University of Nairobi herbarium and a voucher specimen deposited therein. The plant materials were air dried for a week before being milled into powder. The plant was confirmed at the University of Nairobi herbarium and a voucher specimen deposited. The powder (1 kg) was macerated in distilled water in a weight: volume ratio of 1:8 for twenty (20) minutes and made to 8 litres of solution. The resulting suspension was then filtered. Filtration of the solution was first done using cotton wool and then Whatman’s filter paper. The resulting filtrate was frozen and then underwent lyophilization done to obtain a freeze-dried extract. The freeze-dried extract was weighed, placed in amber coloured sample bottles and stored in a deep freezer.

### Animal experiments

Twenty freshly weaned Sprague Dawley (SD) rats weighing around 95 g were used in the experiment. They were housed, 5 members each, in metallic cages (dimensions 109 cm by 69 cm by 77.5 cm) with floor covered with wood shavings. The shavings were replaced thrice every week. Lighting of the cages was maintained at a 12-hour day and night cycle. For the first 8 weeks, all the animals were fed on a high fat (15%) and high fructose (20%) diet *ad libitum.* Weight and Fasting (5 hour fast) blood glucose levels including oral glucose tolerance tests were measured on both day 0 and day 56 (last day of 8^th^ week). On day 1, the animals were confirmed to be non-diabetic as the FBG levels were on average 4.38 +/- 0.33 mmol/l which was less than 7.5 mmol/L, a limit suggested by Wang *et al*[4]. The blood drawn was hence labelled as non-obese (Nob) and stored. The weights and blood glucose level values (averaged 325 g and 6 mmol/L respectively) obtained on the 56^th^ day were used to designate the rats as obese. The rats were then regarded as obese (with metabolic syndrome). These animals were thereafter randomly grouped, with 5 members each, into: Obese (Ob; fed on high fat 15%, high fructose 20% diet *ad libitum* as before, intraperitoneal injection of Normal Saline: 0.9% NaCl in water), Oxytocin treated (Oxy; same feeding as obese and intraperitoneal injection of oxytocin 1 mg/kg), *Uvariodendron anisatum Verdeck (Annonaceae)* (UAV) extract treated at low dose (LDOx; same feeding as obese and intraperitoneal injection of low dose of UAV at 100 mg/kg of body weight) and high dose (HDOx; same feeding as obese and intraperitoneal injection of low dose of UAV at 200 mg/kg of body weight). The oxytocin and UAV treatment were carried out for 7 days. The solvent used in dissolving both the oxytocin powder (Sigma-Aldrich, USA) and the freeze-dried UAV extracts was normal saline (0.9% NaCl in water). All the prepared solutions of oxytocin and the extracts were administered daily on a 24-hour interval. Blood glucose testing was done using a commercial glucometer (StatStrip Xpress Nova Biomedical, Waltham MA, USA) and weight measurement using an electronic beam balance. The blood samples (∼50 µL) were drawn from each rat via lateral vein sampling after local anaesthesia of the tail by topical application of Lidocaine. All the rats were then euthanized following an overnight fast using 20% Phentobarbital (1ml/kg of body weight) injected intraperitoneally at the end of the experiment. Confirmation of death was via loss of the pupillary light reflex. The drawn blood from each rat was stored in sodium citrate vacutainers to prevent clotting and refrigerated at 4°C.

### Raman spectroscopy

Raman spectroscopy was carried out using confocal Raman system (STR, Seki Technotron Corp) equipped with a 785 nm laser and a spectrometer (Princeton Instruments). The conductive silver paste smeared microscope glass slides used as Raman sample substrates were prepared as described in [25]. Spectral callibration of the Raman spectroscopic device was also done as described in reference [25]. The experimental parameters for this study were as follows: grating, 600 groves/mm; Centre wavelength, 850.97 nm (980 cm^−1^); excitation power at sample position, ∼9 mW; spot size at sample position, ∼65 µm; exposure time, 10 sec; spectral accumulation, 5 sec; X10 microscope objective (Olympus MPlanFN, 0.3NA). A small amount of blood (∼ 10 µL, whole blood) was pipetted onto the silver smeared glass slide and air dried for 1 hour. Ten spectra at ten random spots per rat’s blood sample were recorded making a total of 250 (50 data sets for non-obese samples included) spectral data sets with each group having 50 data sets.

### Data processing

Data pre-processing involved smoothing using Vancouver algorithm based on fifth-order polynomial fitting method developed by Zhao *et al*[26]. Then followed by series of normalizations: to maximum intensity, to area under curve and subtraction of minimum value per data set in that order in MATLAB 2017a scripting environment. For plotting, ORIGIN (Origin pro 9.1) software was used.

### Spectral similarity

The spectral similarity between data from UAV freeze dried extracts and those from oxytocin powder and solutions in normal saline was computed using Cosine similarity. Cosine similarity is defined as[27,28]; 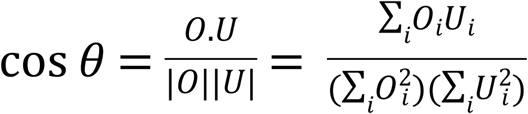. Where *O*_*i*_ and *U*_*i*_ denotes Raman intensity at wavenumber i of spectra from oxytocin and from UAV extract respectively. Identical (dissimilar) spectral data sets will give a value of 1 (0) while directly opposite data sets gives a value of −1. A positive value of cos *θ* close to 1 implies that the two Raman spectral data sets are identical.

### Principal component analysis

Principal component analysis (PCA) was used in segregating between the Raman spectral data sets from blood of non-obese, obese and the treated SD rats. This analytical technique utilizes spectral patterns in the differentiation. The combined data set of all the samples was used. The spectral pattern variations are expressed in terms of percentage variance and ranking done[29,42]. The results are represented on a set of orthogonal axes referred as principal components (PCs). The PC with the highest variance is called PC1, flowed by PC2 and so on [37]. Each of the spectral data set is displayed as a point (score) on a PC plane. In this work a score plot of the first two PCs with the highest variance was done.

### Ethical approval

Ethical approval for the study was granted by the Biosafety, Animal Care and Use Committee, Faculty of Veterinary Physiology, University of Nairobi.

## 2. Results and Discussion

### 2.1 Raman spectra of oxytocin and freeze-dried extracts of Uvariodendron anisatum Verdeck (Annonaceae) (UAV)

The Raman spectra from UAV freeze dried extract’s powder and those from oxytocin solution in normal saline displayed identical profiles and so indicating similar Raman active molecular composition or similar bonds (see Fig. 1a). The cosine similarity value obtained from the two was 0.95 indicating that indeed they were almost the same. The cosine similarity value obtained from comparing Raman spectra of extract’s solution and that from oxytocin powder was 0.57 showing that they were not that similar as seen in Figure 1b. The exact reason why the Raman spectra from extract’s powder and oxytocin solution (extract’s solution and that from oxytocin powder) were identical (dissimilar) was not known. It was thought that the interaction between the silver smear and the samples are responsible for the observed variations. The interactions must have influenced conformations of the various bonds in the oxytocin hormone (which is a nanopeptide) [14] and causing the disulphide S-S stretching mode at 508 cm^−1^ in the oxytocin powder to be red-shifted to 401 cm^−1^ in the extract’s solution [14,34]. These same signals were observed to be broad in Figure 1a. The conformational angle must have been less than 60° about the C-S bond as was argued earlier by Maxfield and Scheraga [14]. The other prominent bands observed were those centered at wavenumbers 645 cm^−1^, 846 cm^−1^ and 1617 cm^−1^ ascribed to tyrosine with the commonly known 830/850 cm^−1^ doublet seen in oxytocin powder and solution [34,35]; 1240 cm^−1^ assigned to amide III with anti-parallel β-sheet structure [34]; 1450 cm^−1^ assigned to C-H deformation in isoleucine and 1658 cm^−1^ attributed to amide I vibrations with anti-parallel β-sheet conformation [34].

**Figure 1.**
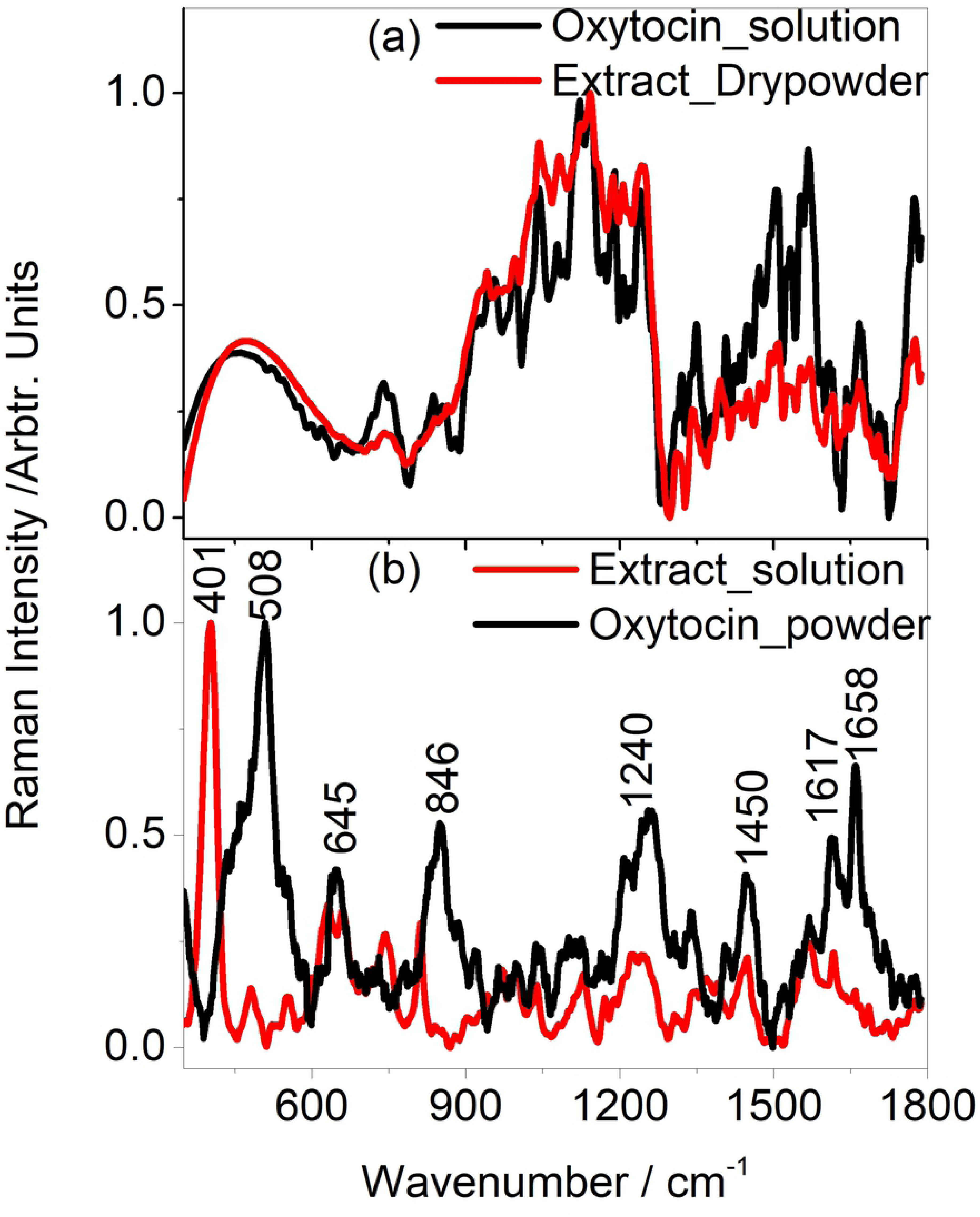
Figure displaying Raman spectra obtained from (a) UAV extract’s dry powder and oxytocin solution and (b) UAV extract’s solution and oxytocin powder. All the samples were placed on conductive silver paste smeared glass slides.

### 2.2 Raman spectra of blood from SD rats

The Raman spectra of blood obtained from SD rats that were obese (Ob), non-obese (NOb), obese and administered oxytocin (Oxy), obese and administered UAV’s extract at low dose (LDOx) and high dose (HDOx) are displayed in Figure 2a. The intense peak at 401 cm^−1^ also seen in extract’s solution (see Fig. 1b) was present in all blood from rats administered oxytocin and UAV extracts but less significant in obese and non-obese rats. This band may be used as oxytocin biomarker band in blood and reflects elevated levels of the hormone in the treated animals. In other murine studies, subjects administered oxytocin exhibited increased levels of the hormone (i.e. oxytocin) in serum [10] and in plasma [7] thus supporting our observation through the assigned Raman peak. Elsewhere, it was reported that in human subjects that were obese and with type II diabetes mellitus, levels of oxytocin were significantly lower compared to healthy subjects [8,19]. The band around 478 cm^−1^ was associated with both fructose and glucose’s skeletal vibrations with tentative assignments; C-C-C, C-C-O, C-O deformations and C-C torsional vibrations [36]. The bands at 638, 812 and 1217 cm^−1^ were attributed to fructose and tentatively assigned to ring deformation and C-C stretching vibrations respectively [36]. Interestingly, the bands at 638 and 812 cm^−1^ (fructose bands) exhibited a decrease in intensity upon administration of both oxytocin and UAV extracts to the diabetic rats as seen in Figures 2b and 2c. In order to interpret this trend, solutions of fructose in normal saline were prepared with concentrations ranging from 0.005 – 0.015 mMol/L and Raman spectra obtained after pipetting onto the conductive silver coated glass slides. The trend of the average intensity of peak around 812 cm^−1^ as a function of fructose concentration (see Figure 2d) was identical to that from blood of SD rats (Figure 2b and 2c). The implication of this was that fructose levels in blood of treated rats (oxytocin- and UAV extract-treated) were higher than in obese and non-obese rats. This trend comparison also indicates that obese rats have the lowest blood fructose concentration as compared with non-obese animals. At the same time, oral administration of oxytocin and UAV extracts causes elevated levels of circulating fructose in SD rats.

**Figure 2:**
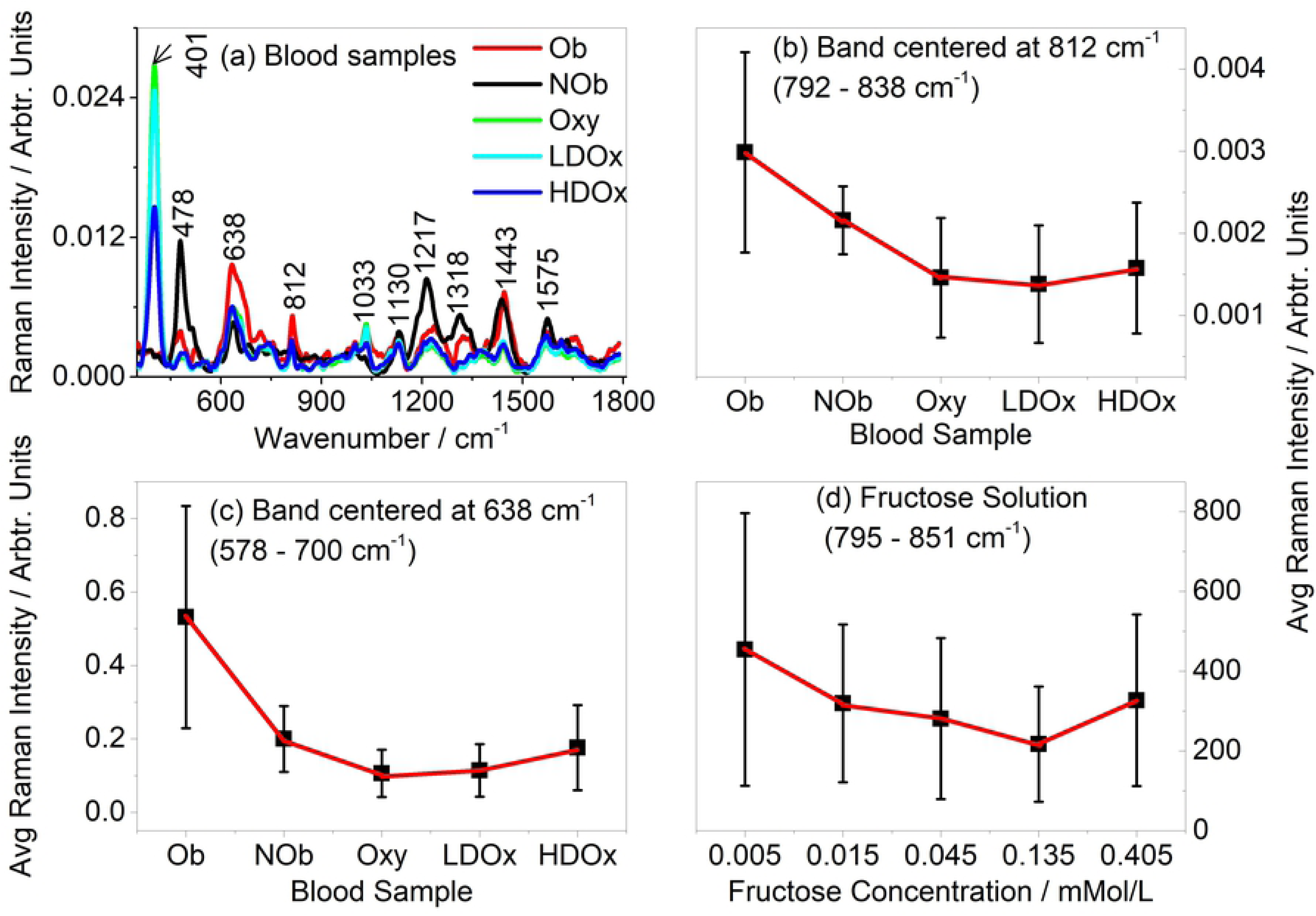
Figure showing (a) Average Raman spectra from blood of obese(Ob), non-obese (NOb), oxytocin treated obese rats (Oxy) and UAV extracts treated obese rats at low dose and high dose (LDOx and HDOx) SD rats (b) and (c) Average Raman intensity of peak centered at 812 cm^−1^ and 638 cm^−1^ respectively and (d) Average Raman intensity of peak centered around 812 cm^−1^ from fructose solution in normal saline.

This Raman spectroscopic study indicates an upregulation of fructose metabolism in the liver [37–39] of SD rats administered oxytocin and UA extracts. This would explain the implied increased concentration of fructose in blood of these animals. It should also be noted that during treatment, the animals were still on a high fat and high fructose diet. The high levels of fructose in blood are usually filtered out through the kidneys and it is expected that their levels in urine are high as reported elsewhere in diabetic humans[40]. In other studies, intraperitoneally injected oxytocin on mice resulted in reduced fructose concentration in seminal vesicles and coagulating glands [20]. The work here, therefore, suggests that intensity of Raman spectral bands assigned to fructose at 638 and 812 cm^−1^ could be used as biomarkers of obesity based on fructose level variation. That is, whenever the intensities of these bands increase above a certain set level per band, then it would mean that the subjects are becoming obese. This would also require reproducible data pre-processing techniques. The bands can also be useful in comparative study between conventionally used anti-obesity medicines and similarly used traditional plant extracts. The other bands at 1033, 1130, 1318 and 1443 cm^−1^ are associated with the branched chain amino acids (BCAAs). The peaks around 1033 and 1130 cm^−1^ are ascribed to C-N stretch, NH_3_ rocking, HCCH torsional vibrations in leucine [41]; CO stretch, OH bending vibrations in both valine and isoleucine [41].

### 2.3 Principal component analysis (PCA)

The Raman spectral data from blood of obese (red squares) were clearly differentiated from those of non-obese (black squares) rats as seen in the PCA score plot of Figure 3a. The segregation between the rats administered oxytocin and UAV extracts was not clear which meant that their Raman spectral profiles were almost identical due to presence of similar Raman active molecules. Through the loadings plot of Figure 3(b,c), PCA shows that the levels of fructose (see peaks at 638, 812 and 1217 cm^−1^) and BCAAs (peaks at 478, 1318 and 1443 cm^−1^) in blood get modified as the bands associated with them play a big role in spectral data segregation. The result further confirms the argument that concentration of fructose in blood of obese rats are different from those of non-obese and treated animals.

**Figure 3:**
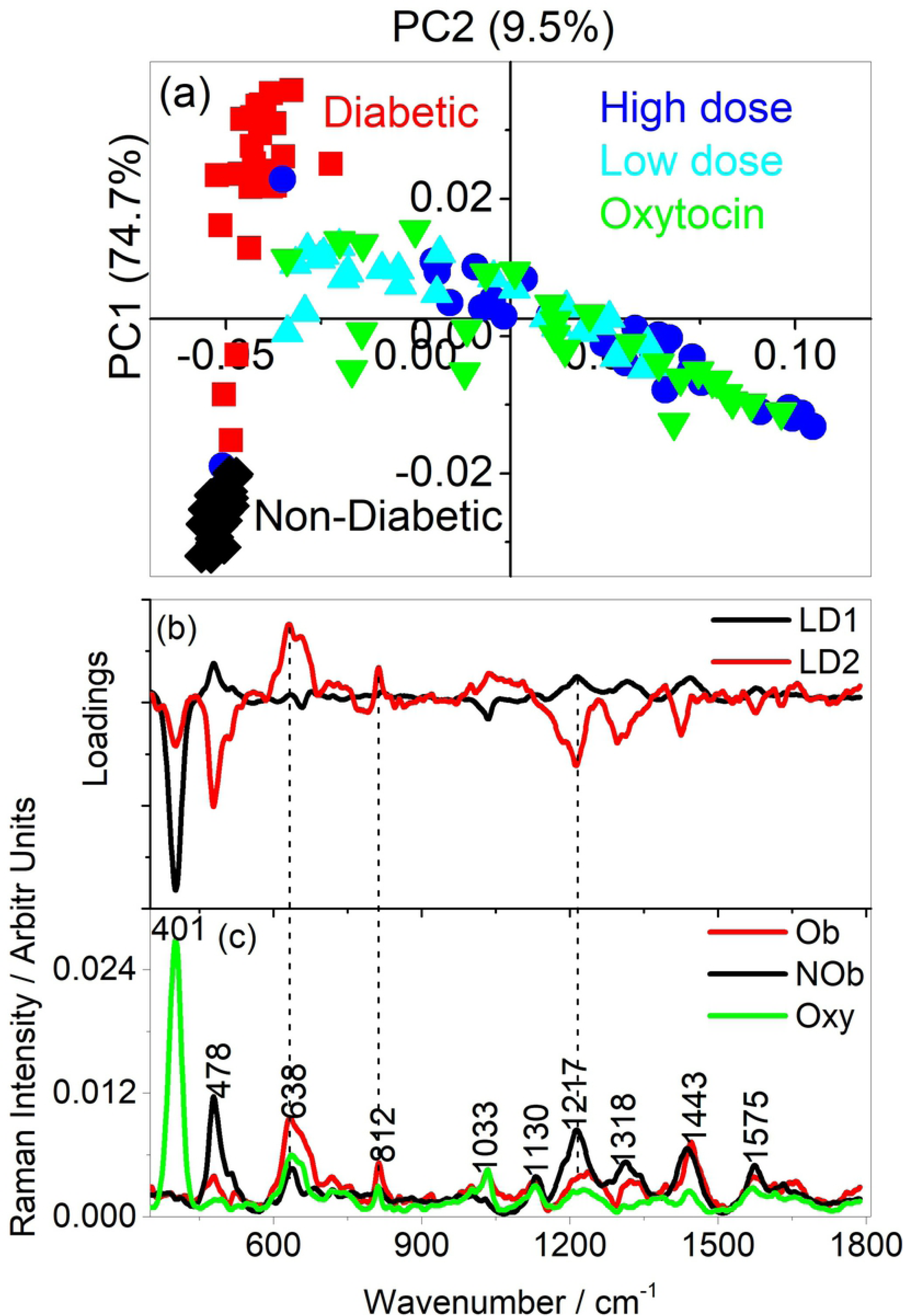
Figure displaying PCA (a) score plot (b) loadings plot for PC1 (LD1) and PC2 (LD2) and (c) Raman spectra from blood of obese, non-obese and oxytocin treated rats. PC1 and PC2 had explained variance of 74.7% and 9.5% respectively. The fructose bands at 638, 812 and 1217 cm^−1^ are indicated with dotted lines and are seen to have played a big role in the segregation of data by PCA as seen in the loadings plot.

The results of the study indicate that Raman spectroscopy can be used as a label free obesity detector or screener with bands associated with fructose as biomarkers. At the same time, the results show that Raman profiles from blood of oxytocin treated rats and those treated with UAV extracts contained similar Raman active molecules. The two compounds (i.e. oxytocin and UAV extracts) influenced obesity in the rats since the spectral profiles were modified. This was also supported by the fact that the average weights and FBG values of the treated animals decreased (325 g to 260 g and 6.3 +/- 0.3 mmol/L to 4.7 +/- 0.4 mmol/L respectively) in the first 7 days after commencing treatment. The low and high doses of the UAV extract did not exhibit discernible differences on the rats as the blood had identical profiles. Herbalists and traditional birth attendants in parts of rural Kenya use UAV to induce labor [23,24]. The Raman study results reveals that the herb has identical effects in obese SD rats as the conventional oxytocin. This implies that the herb is composed of similar Raman active molecules. Further studies need to be done to validate the Raman spectroscopic results reported here as this, to the best of our knowledge, is the first such investigation as regards comparison of UAV and conventional oxytocin and on their influence on obesity in rats.

## 3. Conclusion

The study revealed that Raman spectroscopy can be a powerful tool for detecting obesity with bands associated with fructose acting as biomarkers. It also showed that the technique can be used in performing comparative study of influence of anti-obesity drugs on animal models. Apart from fructose levels in blood getting modified in obese animals, branched chain amino acids are influenced as well. The method further showed that oxytocin and UAV had similar influence on the obese rats and so indicating that both are composed of identical Raman active molecules.

